# Does diet influence physiological and behavioural responses to predation risk?

**DOI:** 10.64898/2026.07.28.741215

**Authors:** Adeeb M. Kunjali, Maria Thaker

## Abstract

As one of the strongest selective forces, predation risk induces antipredator responses in prey that not only maximise survival probability but are energetically expensive. While these responses are known to affect foraging patterns and diet composition, the functional role of changes in dietary macronutrients remains unclear. We investigated the effect of dietary NPE (non-protein energy) concentration on the antipredator responses of the Indian rock agama *Psammophilus dorsalis*. In a manipulative experiment, wild-caught lizards were fed either a low or high NPE diet and exposed to repeated simulated predator attacks. We measured the space-use and hiding duration of lizards after each attack, as well as their circulating corticosterone concentrations at the end of the experiment. Predator exposure induced refuge use for all lizards. Diet had a context-dependent effect: lizards on the high NPE diet used riskier perch locations more frequently than those on low NPE diets, but only during safe periods and not during active threat. Physiologically, lizards on the low NPE diet had significantly higher baseline corticosterone levels, indicating a greater allostatic load. Overall, while dietary NPE did not alter reactive escape behaviour, it influenced the underlying physiological state and proactive behavioural decisions.

## Introduction

Predation risk is one of the strongest selective forces, shaping adaptations and inducing responses in prey that maximise survival probability. These changes, however, are energetically costly. For many animals, an encounter with a predator elicits an immediate escape response, which is followed by behaviours that minimise the risk of further encounters [1]. These behaviours include an increase in refuge use [2–4], avoidance of risky areas [5], or shifts in activity patterns [1,5,6]. For example, some prey reduce overall activity to minimize detection [7], while others increase activity to facilitate escape [8]. The specific behavioural strategy employed is a calculated decision based on an individual’s assessment of vulnerability and the immediate risk [1]. If prolonged, antipredator behavioural strategies to minimise detection by predators can result in missed opportunity costs [9–11], such as that typified by the food-safety trade-off [12]. By avoiding risky times and risky areas, foraging opportunities for prey are constrained, resulting in altered diets and an overall reduction in food acquisition [4,13,14].

Predation risk also elicits physiological responses in prey. In response to predator encounters, most vertebrates increase the concentration of circulating glucocorticoid hormones [15,16]. Glucocorticoids, such as corticosterone, facilitate immediate processes that promote survival, by working on multiple pathways simultaneously [17–19]. During this emergency response phase, corticosterone synergizes with catecholamines to break down glycogen stores and sustain glucose availability through gluconeogenesis [17–19].

Glucocorticoids also inhibit glucose transport in peripheral tissues, thereby sparing circulating glucose for essential organs like the brain [17, 18]. This inhibition of peripheral glucose utilization, combined with sustained gluconeogenesis, extends the metabolic stress response for several hours after the initial catecholamine surge subsides [17]. In addition to these metabolic effects, corticosterone plays a role in facilitating escape responses [20], sharpening cognitive functions and facilitating memory consolidation, all of which have behavioural consequences that are critical for predator avoidance [18–22]. During predation risk, elevated glucocorticoid levels also suppress activities not essential for survival, such as growth and reproduction [17,18,23–26]. The magnitude and longevity of these behavioural and physiological responses to predation risk depends on the intensity, predictability, and frequency of predator encounters [19,27,28].

Ultimately, behavioural and physiological responses to predation risk are energy intensive. According to the general stress paradigm [14], elevated metabolic rates during stressful periods result in a higher demand for energy [17–19]. Behavioural escape responses that require rapid locomotion or hiding without an opportunity to forage also require additional energy. In many taxa, this additional energy requirement is obtained by increasing glycogenolysis and by increasing foraging effort to rebuild depleted energy stores [13,14,29,30]. Increase in feeding behaviour is mechanistically mediated by glucocorticoids that act on the brain to increase appetite and feeding behaviour, especially promoting carbohydrate intake once the immediate threat has passed [17, 19]. Having access to foraging options that vary in nutritional content could therefore influence subsequent antipredator responses. While several studies have found that prey shift their diets to higher carbohydrate foods in response to the stress of predation risk [31,32], it is unclear whether differences in dietary carbohydrate and lipid levels subsequently modulates prey responses to predators.

In this study, we investigate the effect of dietary non-protein energy (NPE; carbohydrate and lipid) concentration on the antipredator responses of an ectothermic vertebrate. Using the wild-caught Indian rock agama, *Psammophilus dorsalis*, as a model system, we conducted a manipulative experiment wherein lizards were provided with either a low carbohydrate+lipid or a high carbohydrate+lipid diet. These lizards were exposed to both safe conditions, where they were left undisturbed, and risky conditions, where they were repeatedly attacked by a simulated predator. We measured overall space-use and hiding duration during both these conditions, as well as baseline corticosterone levels at the end of the experiment. If NPE fuels antipredator responses, we predict that lizards on a high-NPE diet will have bolder space use characterised by more time exposed and not in refuge, shorter hiding duration after attacks, and lower baseline corticosterone levels. Our study provides the missing link between dietary shifts and antipredator responses, by specifically testing the functional benefit of consuming higher carbohydrates and lipids when under predation risk.

## Materials and methods

Adult males of *Psammophilus dorsalis* (Figure 1(a)) lizards (N = 47) were captured from northern Bangalore (13° 11 ’08.3”N 77° 36’ 22.3”E) between August and October 2023, towards the end of the breeding season for the species. Lizards were caught by lassoing and were transferred to the field station within four hours of capture, where their snout-vent length (SVL range: 108-145 mm) and mass (mass range: 45-109 g) were measured. Lizards were then housed in separate semi-outdoor enclosures (4.9 m long × 1.4 m wide × 1.2 m high) that had a soil substrate surrounded by foam walls and contained a refuge comprising a clay shingle on cement bricks, two cinder block perches (one near the refuge and one at the opposite end), and water-filled petri dishes at each perch (Figure 1(b)). Adjustable openings were made to allow for the introduction of food and to enable maintenance and cleaning of the enclosure. A CCTV camera was placed above each enclosure and was set to continuously record the lizards from 8:00 to 16:00 hours throughout the experiment. Eight of these enclosures were situated in a larger tiled-roof structure with mesh walls that prevented rain and natural predators from entering the enclosures. This entire set-up was outdoors and so lizards were exposed to natural light and temperature conditions, including sunlight for basking. On the day of capture, lizards were provided with water but not food and were allowed to acclimate to the enclosure for 24 hours before the start of the experiment. The total time in captivity was 8 days, after which all individuals were marked with non-toxic ink on their ventral side and released at their site of capture.

**Figure 1:**
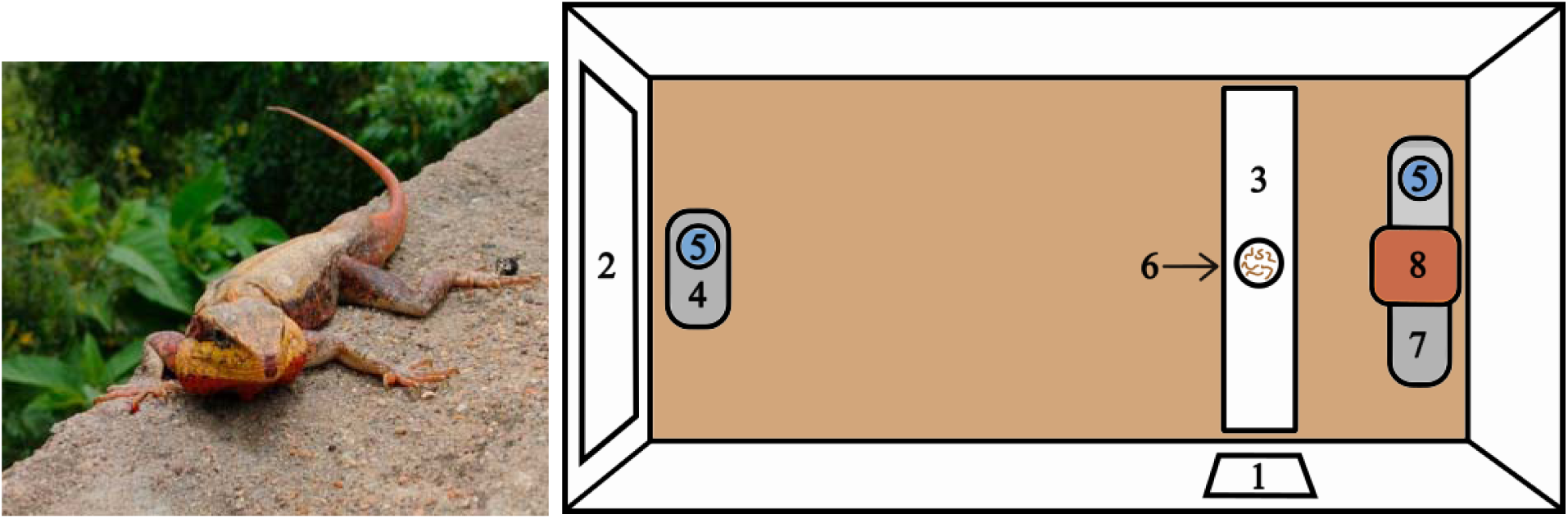
Male of *Psammophilus dorsalis,* and schematic of experiment enclosure, labelled with the openings (1) to introduce food and (2) allow the researcher to enter, (3) white foam board on which mealworms (food) was presented, (4) perch far from the refuge, (5) Petri dish containing water (one at each perch), (6) Petri dish containing food, (7) Perch near the refuge, and (8) Refuge.

### Diet Treatment

Lizards were randomly assigned to either a low NPE (N = 23) or high NPE (N = 24) diet, which they received throughout the experiment. The two diets were prepared by rearing mealworms for at least 10 days in either oats (low NPE diet) or in 100% Maltodextrin based complex carbohydrate powder (high NPE diet). High-NPE mealworms were additionally coated with carbohydrate powder before they were provided to lizards.

To analyse the macronutrient composition of diets, we first dried and crushed mealworms from each diet type and ran the following analyses in triplicate. Carbohydrate content in these crushed samples were quantified using the anthrone method after NaOH extraction, and protein content was quantified with a colorimetric assay following the Bradford method using a kit (TP0100, Sigma-Aldrich) [as per 33,34]. We also quantified the amount of unsaturated lipids in the mealworms using a colorimetric phospho-vanillin assay (modified from [33]). This method quantifies only the macronutrient content of the soft tissues of mealworms and does not account for macronutrients embedded within the chitinous exoskeleton that is indigestible to predators [33].

Mealworms in the low NPE diets contained 0.49 ± 0.06 SE mg carbohydrate, 22 ± 7 mg lipids, and 26.08 ± 1.58 mg protein per 100 mg mealworm. Mealworms in the high NPE diets contained 3.15 ± 0.33 SE mg carbohydrate, 35.2 ± 9.7 mg lipids, and 28.44 ± 1.10 protein per 100 mg mealworm.

### Experimental Protocol

The experiment consisted of three phases: habituation (days 1–2), safe (days 3–4), and predator exposure (days 5–6). During the habituation and safe phases, lizards were provided with 15 mealworms (of either high NPE or low NPE types) every day between 1030 and 1130, coinciding with peak lizard activity. The weight of mealworms was measured before and after every feeding period to determine the amount of food eaten by each lizard everyday. One individual that did not eat during the habituation phase was excluded from the study. Apart from food provision, lizards were left undisturbed during the habituation and safe phases.

On days 5 and 6, we simulated predator attacks using an eagle model attached to a pole (Figure S1). Each attack, which involved rapidly approaching the lizard with the eagle model, lasted sixty seconds and always induced escape behaviour. Six attacks were performed daily: three before feeding (0800–1000) and three after (1300–1500), with each attack at least an hour after the last one.

### Behavioural Measures

As a measure of spatial activity, we used the CCTV video recordings and quantified time spent in three spatial zones of the enclosure: within the refuge, on the perch near the refuge, and on the perch far from the refuge. We converted these daily duration values to the proportion of time spent in each zone by dividing them against the total recording time for each phase (safe and predator exposure). In response to predator attacks, we also recorded the time it took to emerge from the refuge after each attack, which we denote as hiding duration.

### Corticosterone Measures

At the end of the experiment (day 7), we collected blood samples from the retro-orbital sinus of all lizards using heparinized microcapillary tubes (following methods in [2]). Lizards were caught between 0600 - 0700 and a blood sample (∼100 ul) was obtained within 4 minutes of capture. Samples were centrifuged and the plasma was stored at −20°C until hormone analysis. To avoid the stress of repeated blood sampling, only a single sample was taken at the end of the experiment, as opposed to during both the safe and predator exposure phases. We measured corticosterone levels using Arbor Assay DetectX kits (CORT K014-H) that have been optimised for this species. We analysed 4µl of plasma at a dilution ratio of 1:100 in triplicate. The intra-assay coefficient of variation was 0.46 - 15.2% and the inter-assay coefficient of variation was 8.56%.

### Statistical Analyses

Food intake data (weight of mealworms consumed) deviated from the assumptions of normality across all groups (Shapiro-Wilk test, p < 0.001 for all diet × phase combinations). We first report the incidences of non-feeding (0 mealworms consumed) and then for those that did eat at least 1 mealworm, we determined differences in food intake using a GLMM (gamma distribution and log link function) with diet (low or high NPE), phase (safe or predator exposure), and their interaction as predictors. We included lizard ID as a random effect as the same lizard was tested across all phases.

The proportion of time spent in each part of the enclosure (W = 0.71–0.97, p < 0.02), hiding duration (W = 0.95, p < 0.05), and corticosterone concentrations (W = 0.90, p < 0.05) were also non-normally distributed as assessed by the Shapiro-Wilk test. We determined differences in time allocation in different parts of the enclosure using a GLMM (beta family with logit link function), with diet (low or high NPE), phase (safe or predator exposure), and their interaction as predictors. We applied a Smithson-Verkuilen transformation to proportions containing zeros (time in refuge: 3.2% zeros; time on far perch: 12.8% zeros), and included lizard ID as a random effect. To determine differences in hiding duration after a predator attack, we used a GLMM (gamma distribution and log link function) with attack number (1–6), diet (low or high NPE), and an interaction between attack number and diet as predictors. Corticosterone levels at the end of the experiment were analysed with a GLM with a gamma distribution and log link function, with SVL and diet as fixed predictors. Odds ratios were calculated and reported wherever relevant.

## Results

### Food consumption

There were 15 instances (7.98%) of non-feeding out of 188 feeding opportunities, of which 12.77% (n=12) were in the predator exposure phase and 3.19% (n=3) were in the safe phase. Diet type did not affect the incidence of feeding as 8.70% (n = 8) of feeding opportunities for the low NPE diet and 7.29% (n=7) for the high NPE diet were missed. For those lizards that did eat, the amount of mealworms (g) consumed by lizards was not significantly affected by phase (β = −0.11, Z = −1.62, p = 0.105), diet (β = −0.01, Z = −0.11, p = 0.910), or their interaction (β = 0.10, Z = 1.02, p = 0.306, Figure 2; Table S1).

**Figure 2:**
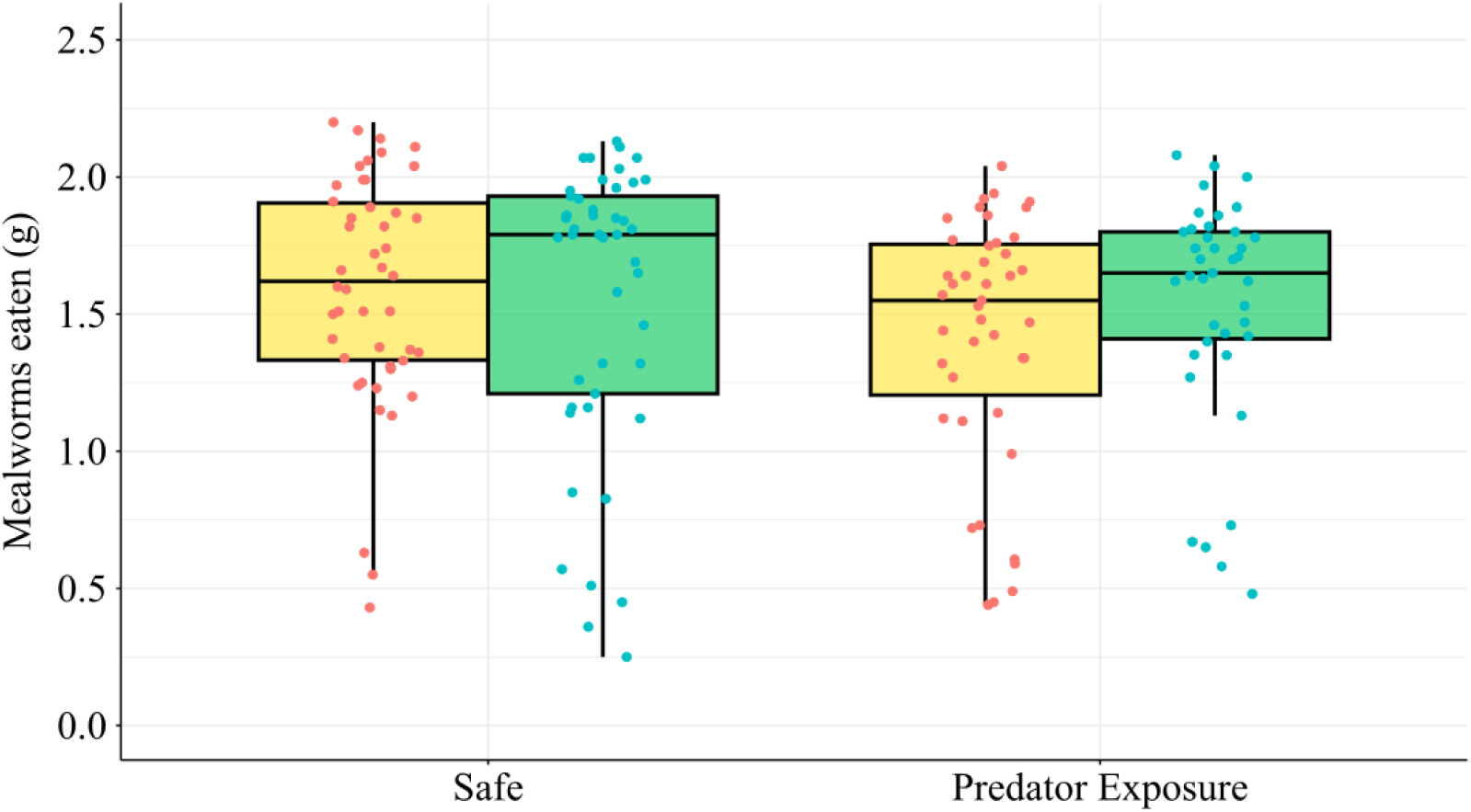
Mealworms eaten (mass in g) by lizards from the high NPE (yellow) and low NPE (green) diets during the habituation, safe and predator exposure phases. Shown are boxplots with interquartile ranges. Actual values are plotted as red and blue points.

### Space-use

The behaviours of lizards were significantly influenced by whether animals were in the safe phase or exposed to predation risk. Lizards increased their refuge use (β = 1.76, Z = 8.44, p < 0.001, Figure 3a, Table S2) and decreased use of the perch near the refuge (β = −0.87, Z = - 5.67, p < 0.001, Figure 3b, Table S3) during the predator exposure phase compared to the safe phase. There was no significant effect of diet or an interaction between diet and phase on the use of the refuge or the perch near the refuge (Tables S2, S3).

**Figure 3:**
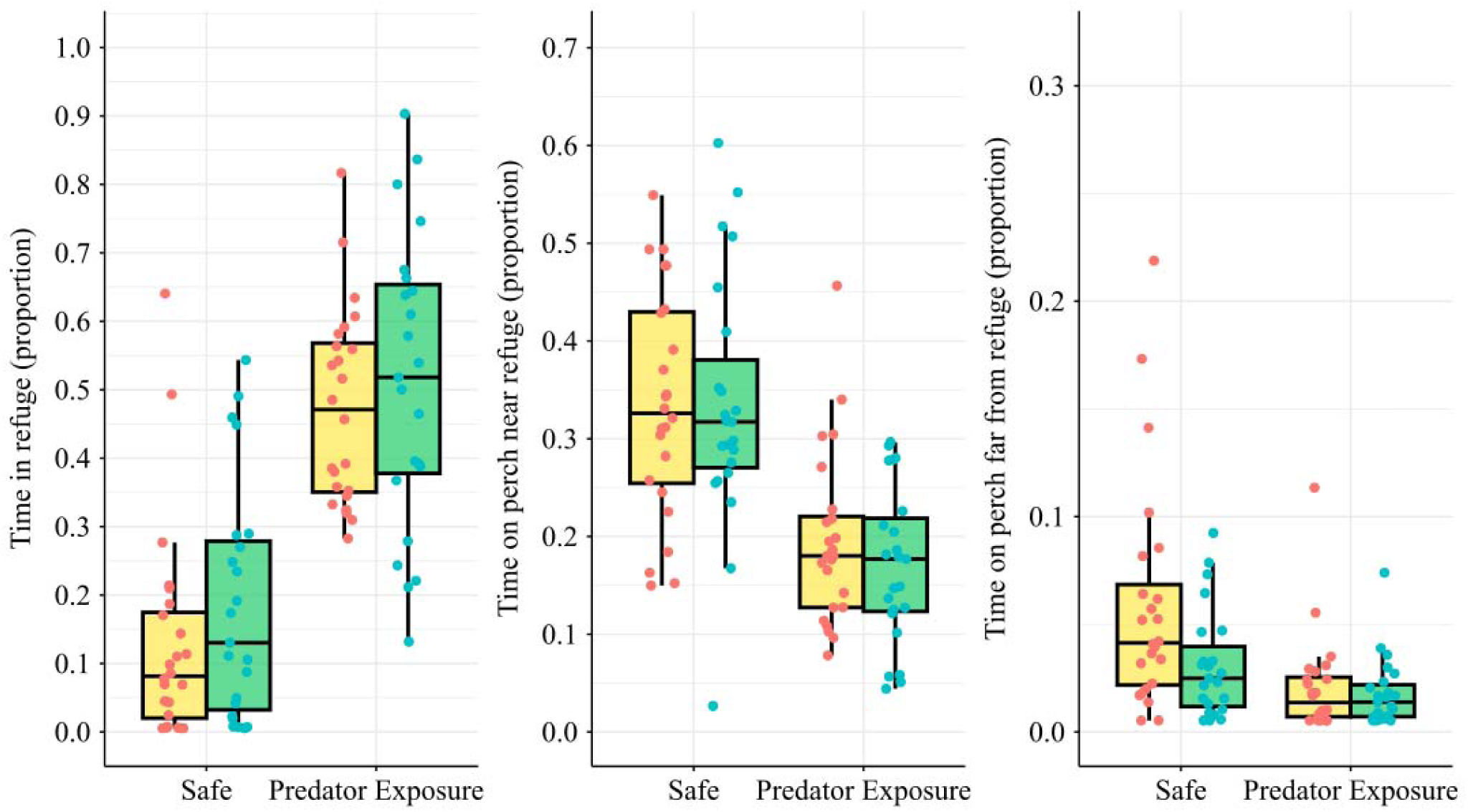
Proportion of time spent (a) in refuge, (b) on the perch near the refuge, and (c) on the perch far from the refuge by lizards from the high NPE (yellow) and low NPE (green) diets during the safe and predator exposure phases. Shown are boxplots with interquartile ranges. Actual values are plotted as red and blue points.

There was a significant interaction between diet and phase on the use of the perch far from refuge (β = −0.54, Z = −2.37, p = 0.017, Figure 3c, Table S4). Predator exposure caused lizards from both high NPE (OR = 0.36[0.24, 0.53], Z.ratio = −6.78, p <0.001) and low NPE diet groups (OR = 0.61[0.39, 0.96], Z.ratio = −2.79, p = 0.027) to reduce the usage of the perch far from the refuge compared to the safe phase. During the safe phase, lizards on the high NPE diet had higher perch usage than the lizards on the low NPE diet (OR = 1.83[1.04, 3.21], Z.ratio = 2.76, p= 0.030, Table S5).

### Hiding duration

During the predator exposure phase, predator attacks elicited refuge seeking behaviour in all lizards (Figure 4). Time to emerge from the refuge after an attack was significantly affected by attack number (for attacks 3-6: βs ranged from −0.41 to −0.70, Zs ranged from 3.70 to 6.23, p < 0.001 for all comparisons) but not diet type (β = −0.003, Z = −0.02, p = 0.986) or the interaction between attack number and diet type (p > 0.1 for all comparisons, Figure 4, Table S6). Lizards took longer to emerge from refuge after the first attack compared to the second attack (exp(0.26) ≈ 1.3 times longer, Z.ratio= 3.22, p < 0.016). Subsequently, hiding duration declined with attack number such that attacks 3-6 resulted in significantly shorter emerge times compared to attack 1 and 2 (exp(0.53) ≈ 1.70 to exp(0.72) ≈ 2.05 times shorter than attack 1, Z.ratios ranged from 3.38 to 8.97, p < 0.001 for all comparisons except attack 2 and 3 where p = 0.009, Table S7).

**Figure 4:**
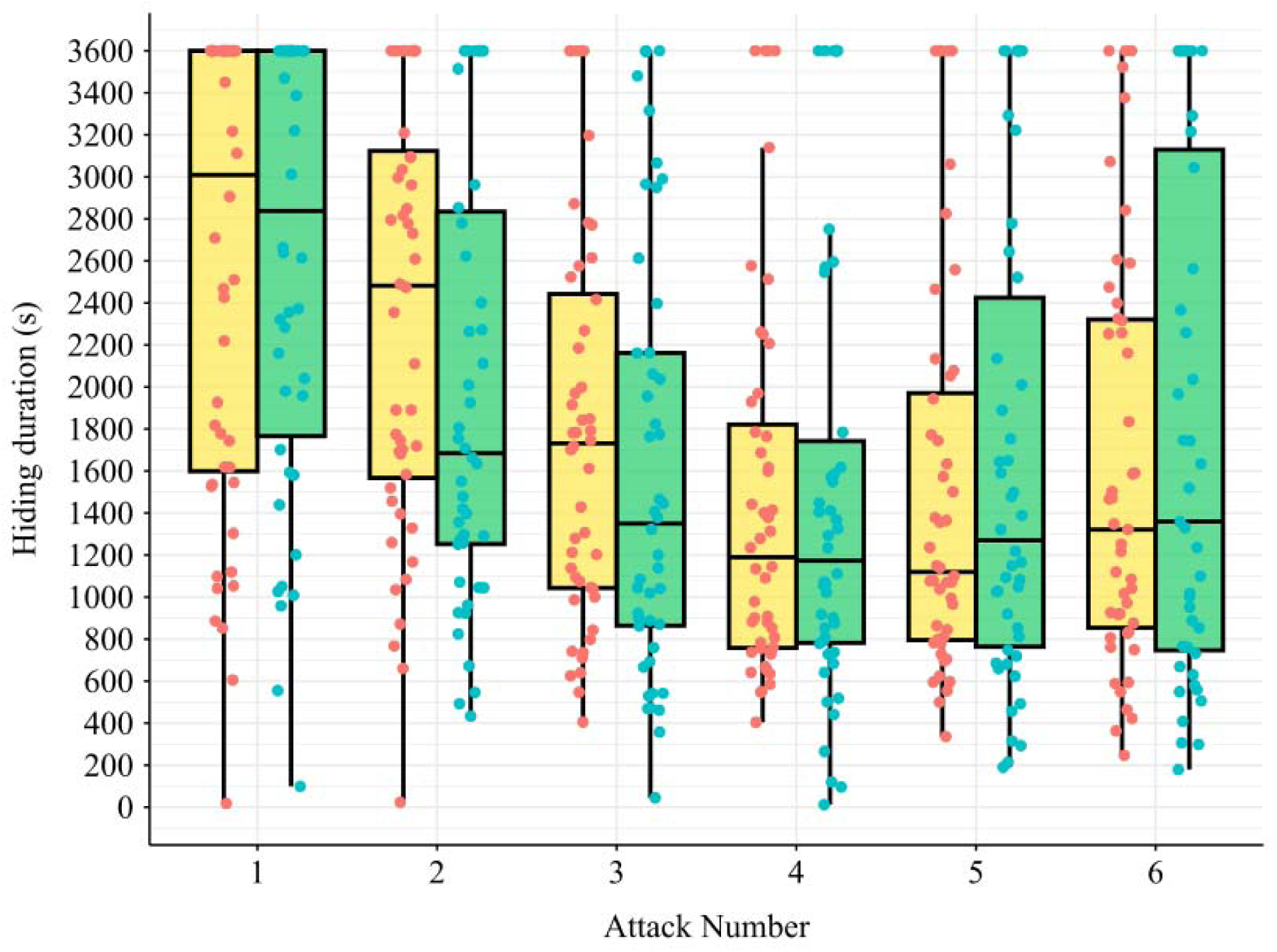
Hiding duration after each predator attack by lizards from the high NPE (yellow) and low NPE (green) diet groups. Shown are boxplots with interquartile ranges. Actual values are plotted as red and blue points.

### Corticosterone level

Baseline corticosterone levels at the end of the experiment, after both safe and predator exposure phases, were significantly affected by diet (β = 0.40, Z = 2.54, p = 0.015). Lizards on low NPE diets had higher levels of baseline corticosterone than those on high NPE diets (Figure 5, Table S8).

**Figure 5:**
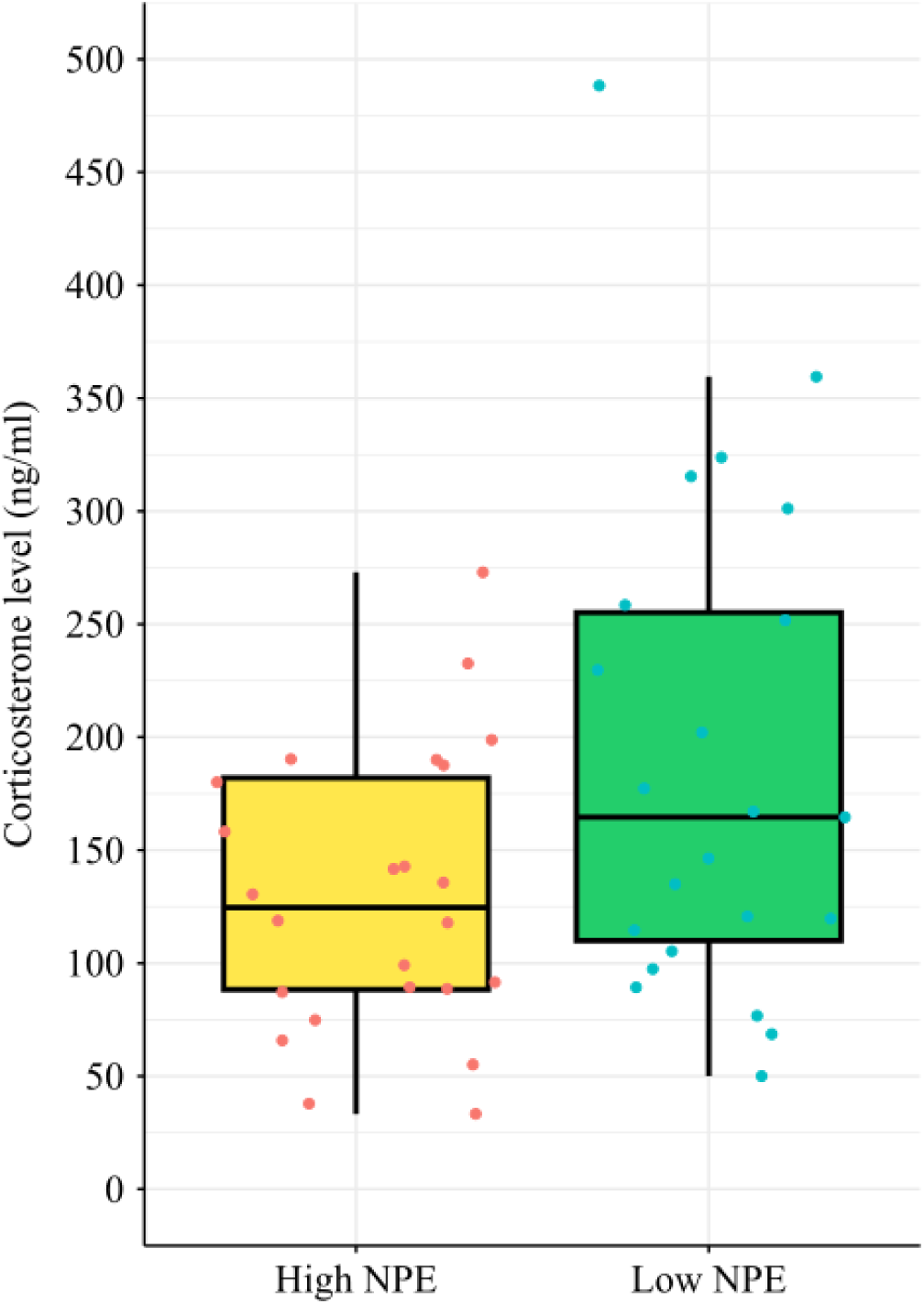
Baseline corticosterone level from the high NPE (yellow) and low NPE (green) diet groups. Shown are boxplots with interquartile ranges. Actual values are plotted as red and blue points.

## Discussion

Predation risk is a powerful ecological pressure, influencing the behaviour and physiology of most animals, including *P. dorsalis*. Animals under predation risk are not only behaviourally constrained from freely foraging [12,20], but are physiologically driven to consume greater carbohydrates [13,31,32]. Here we test whether a shift in diet to higher NPE content modulates antipredator responses. We found that dietary NPE levels had a limited and context-dependent effect on the foraging behaviour and antipredator responses of *P. dorsalis*. The most pronounced effect was on physiology. Lizards on a low NPE diet had higher baseline corticosterone levels at the end of the trials compared to those on a high NPE diet, indicating a potentially higher allostatic load even in the absence of an immediate threat [18,19]. Behaviourally, the high NPE diet did not lead to bolder behaviour during predator exposure as predicted, but instead promoted greater use of risky space during safe conditions. Thus, contrary to our prediction, diet did not seem to influence the immediate behavioural response to an attack or the habituation to repeated attacks.

During the predation risk phase, lizards spent more time in refuge and less time basking out on perches compared to when they were in the safe phase. Consistent with the food-safety trade-off, lizards were also less likely to forage during the predation risk phase compared to the safe phase; although the amount of food eaten by those who foraged was unaffected by predation risk [1]. This suggests that while predation risk influenced the decision to forage, it did not affect how much was consumed once foraging commenced. These risk averse behaviours are consistent with the existing literature on antipredator strategies in other taxa, including lizards [1–5]. During repeated attacks from the predator model, we also found that lizards habituated, as they progressively reduced their hiding duration over time. *P. dorsalis* in the wild also quickly habituate, reducing their flight initiation distance from an approaching human predator by the second “attack” [35].

Predictability and intensity of risk is a key determinant of hiding duration in many animals [36–38]. Given that our simulated predator attacks never resulted in actual physical contact or injury to lizards, it is likely that lizards perceived the threat as relatively low risk. Despite these clear behavioural responses to predation risk, we found that diet type did not significantly modulate the escape response, suggesting that the low NPE diets were sufficient for lizards to maintain their risk averse behaviours, or that they already had resource stores to weather the risky period. Whether dietary state modulates antipredator behaviour under more chronically risky or energetically conflicting scenarios remains to be tested.

Despite its limited effect on behaviour, diet modulated the hormone levels of lizards. Lizards on a low NPE diet had elevated baseline corticosterone, suggesting a higher allostatic load [17–19] than those from the high NPE diet. Behaviourally, the high NPE diet promoted greater use of the perch farthest from refuge only during the safe phase and not during the predation risk phase. While the proactive decision of where to bask and forage in safety was sensitive to the internal energetic state set by diet, the reactive decision to flee and hide from an immediate attack was a more inflexible response that seemed to override the general physiological state of the individual. This pattern aligns with the control of risk hypothesis [39], wherein proactive, flexible responses are used for predictable risk, while reactive, inflexible responses are used for immediate predation risk. However, it has also been shown that some species regulate the allostatic load by modulating corticosteroid-binding globulin (CBG) levels which affect the levels of bioactive free corticosterone [42]. Therefore, it is possible that both changes in corticosterone levels as well as CBG levels, which we did not measure, are involved in managing energy requirements during predation risk. Corticosterone in reptiles also plays a central role in metabolic regulation [17,43], and elevated levels of corticosterone in low NPE lizards could reflect adaptive metabolic adjustment to dietary composition rather than allostatic load alone. Notably, the probability of eating and overall food consumption did not differ significantly in lizards provided with different diet types, and thus the observed responses to predator attacks were not due to differences in food intake, but because of the composition of food.

We acknowledge that the single plasma sample that we obtained provides a snapshot of physiological state at the end of the experiment but may miss the dynamic stress response over time. It is unlikely, however, that diet composition affects the level of corticosterone induced during the predator attacks as physiological responses to acute stressors are more affected by the intensity of the stressor rather than internal state of the prey [39]. Although we know that some prey increase carbohydrate intake and utilization in response to predation risk to meet heightened energy demands, the specific role of non-protein energy in directly or indirectly affecting the antipredator response and survival probability remains unclear and requires further research [14,29–32]. Importantly, for ectotherms, the potential energetic burden of temporary dietary constraints may be mitigated by their capacity for metabolic plasticity [43,44], which likely buffers them against significant energetic costs from short-term challenges like predation-induced diet shifts. Interestingly, we observed a shift in antipredator strategy in some individuals, who transitioned from flight to confronting the model predator during later attacks. This suggests that the behavioural repertoire of species is more complex than a simple flight-hide response [1]. Future studies that explicitly quantify inter-individual variation in antipredator response may reveal state-dependent alternative tactics that influence our understanding of risk-coping strategies.

## Supporting information

Supplementary image and tables

## Acknowledgements

We thank the DBT / Wellcome Trust India Alliance (IA/I/19/2/504639) for funding support to MT. We appreciate the help of S. Ganesh for rearing and maintaining mealworms, and of Avik Banerjee and Mihir Joshi for running the protein, carbohydrate and lipid analyses. We also thank Siddharth Tripathi for helping to score the videos, and Amanda Ben, Mihir Joshi, Pushkar Wagh, and Marwa Abdul Razak for helping to catch lizards from the wild.

## References

1. Lima SL, Dill LM. 1990 Behavioral decisions made under the risk of predation: A review and prospectus. Can. J. Zool. 68, 619–640. (doi:10.1139/z90-092)

2. Thaker M, Lima SL, Hews DK. 2009 Alternative antipredator tactics in tree lizard morphs: Hormonal and behavioural responses to a predator encounter. Anim. Behav. 77, 395–401. (doi:10.1016/j.anbehav.2008.10.014)

3. Avalos A, Cooper, Jr. W. 2010 Predation risk, escape and refuge use by mountain spiny lizards (*Sceloporus jarrovii*). Amphib.-Reptil. 31, 363–373. (doi:10.1163/156853810791769419)

4. Kelleher V, Hunnick L, Sheriff MJ. 2021 Risk-induced foraging behavior in a free-living small mammal depends on the interactive effects of habitat, refuge availability, and predator type. Front. Ecol. Evol. 9, 718887. (doi:10.3389/fevo.2021.718887)

5. Tolon V, Dray S, Loison A, Zeileis A, Fischer C, Baubet E. 2009 Responding to spatial and temporal variations in predation risk: Space use of a game species in a changing landscape of fear. Can. J. Zool. 87, 1129–1137. (doi:10.1139/Z09-101)

6. Ross J, Hearn AJ, Johnson PJ, Macdonald DW. 2013 Activity patterns and temporal avoidance by prey in response to Sunda clouded leopard predation risk. J. Zool. 290, 96–106. (doi:10.1111/jzo.12018)

7. Descalzo E, Tobajas J, Villafuerte R, Mateo R, Ferreras P. 2021 Plasticity in daily activity patterns of a key prey species in the iberian peninsula to reduce predation risk. Wildl. Res. 48, 481–490. (doi:10.1071/WR20156)

8. Li JL, Li HW. 1979 Species specific factors affecting predator prey interactions of the copepod *Acanthocyclops vernalis* with its natural prey. Limnol. Oceanogr. 24, 613– 626. (doi:10.4319/lo.1979.24.4.0613)

9. Orrock JL, Preisser EL, Grabowski JH, Trussell GC. 2013 The cost of safety: Refuges increase the impact of predation risk in aquatic systems. Ecology 94, 573–579. (doi:10.1890/12-0502.1)

10. Abramsky Z, Rosenzweig ML, Subach A. 2002 The costs of apprehensive foraging. Ecology 83, 1330–1340. (doi:10.1890/0012-9658(2002)083%5B1330:TCOAF%5D2.0.CO;2)

11. Eccard JA, Liesenjohann T. 2014 The importance of predation risk and missed opportunity costs for context-dependent foraging patterns. PLoS ONE 9, e94107. (doi:10.1371/journal.pone.0094107)

12. Brown JS, Kotler BP. 2004 Hazardous duty pay and the foraging cost of predation. Ecol. Lett. 7, 999–1014. (doi:10.1111/j.1461-0248.2004.00661.x)

13. Hawlena D, Pérez-Mellado V. 2009 Change your diet or die: Predator-induced shifts in insectivorous lizard feeding ecology. Oecologia 161, 411–419. (doi:10.1007/s00442-009-1375-0)

14. Hawlena D, Schmitz OJ. 2010 Physiological stress as a fundamental mechanism linking predation to ecosystem functioning. Am. Nat. 176, 537–556. (doi:10.1086/656495)

15. Millanes PM, Pérez-Rodríguez L, Rubalcaba JG, Gil D, Jimeno B. 2024 Corticosterone and glucose are correlated and show similar response patterns to temperature and stress in a free-living bird. J. Exp. Biol. 227, jeb246905. (doi:10.1242/jeb.246905)

16. Dantzer B, Fletcher QE, Boonstra R, Sheriff MJ. 2014 Measures of physiological stress: A transparent or opaque window into the status, management and conservation of species? Conserv. Physiol. 2, cou023–cou023. (doi:10.1093/conphys/cou023)

17. Wingfield JC, Maney DL, Breuner CW, Jacobs JD, Lynn S, Ramenofsky M, Richardson RD. 1998 Ecological bases of hormone—behavior interactions: The “emergency life history stage”. Am. Zool. 38, 191–206. (doi:10.1093/icb/38.1.191)

18. Sapolsky RM, Romero LM, Munck AU. 2000 How do glucocorticoids influence stress responses? Integrating permissive, suppressive, stimulatory, and preparative actions*. Endocr. Rev. 21, 55–89. (doi:10.1210/edrv.21.1.0389)

19. McEwen BS, Wingfield JC. 2003 The concept of allostasis in biology and biomedicine. Horm. Behav. 43, 2–15. (doi:10.1016/S0018-506X(02)00024-7)

20. Thaker M, Lima SL, Hews DK. 2009 Acute corticosterone elevation enhances antipredator behaviors in male tree lizard morphs. Horm. Behav. 56, 51–57. (doi:10.1016/j.yhbeh.2009.02.009)

21. Thaker M, Vanak AT, Lima SL, Hews DK. 2010 Stress and aversive learning in a wild vertebrate: The role of corticosterone in mediating escape from a novel stressor. Am. Nat. 175, 50–60. (doi:10.1086/648558)

22. Salehi B, Cordero MI, Sandi C. 2010 Learning under stress: The inverted-U-shape function revisited. Learn. Mem. 17, 522–530. (doi:10.1101/lm.1914110)

23. Silverin B. 1986 Corticosterone-binding proteins and behavioral effects of high plasma levels of corticosterone during the breeding period in the pied flycatcher. Gen. Comp. Endocrinol. 64, 67–74. (doi:10.1016/0016-6480(86)90029-8)

24. Love OP, Breuner CW, Vézina F, Williams TD. 2004 Mediation of a corticosterone-induced reproductive conflict. Horm. Behav. 46, 59–65. (doi:10.1016/j.yhbeh.2004.02.001)

25. Vitousek MN, Jenkins BR, Safran RJ. 2014 Stress and success: Individual differences in the glucocorticoid stress response predict behavior and reproductive success under high predation risk. Horm. Behav. 66, 812–819. (doi:10.1016/j.yhbeh.2014.11.004)

26. Bonier F, Moore IT, Martin PR, Robertson RJ. 2009 The relationship between fitness and baseline glucocorticoids in a passerine bird. Gen. Comp. Endocrinol. 163, 208–213. (doi:10.1016/j.ygcen.2008.12.013)

27. Bennett AM, Longhi JN, Chin EH, Burness G, Kerr LR, Murray DL. 2016 Acute changes in whole body corticosterone in response to perceived predation risk: A mechanism for anti-predator behavior in anurans? Gen. Comp. Endocrinol. 229, 62–66. (doi:10.1016/j.ygcen.2016.02.024)

28. Archard GA, Earley RL, Hanninen AF, Braithwaite VA. 2012 Correlated behaviour and stress physiology in fish exposed to different levels of predation pressure: Stress hormones, behaviour and predation. Funct. Ecol. 26, 637–645. (doi:10.1111/j.1365-2435.2012.01968.x)

29. Teegarden SL, Bale TL. 2008 Effects of stress on dietary preference and intake are dependent on access and stress sensitivity. Physiol. Behav. 93, 713–723. (doi:10.1016/j.physbeh.2007.11.030)

30. Schmitz OJ, Rosenblatt AE, Smylie M. 2016 Temperature dependence of predation stress and the nutritional ecology of a generalist herbivore. Ecology 97, 3119–3130. (doi:10.1002/ecy.1524)

31. Hawlena D, Schmitz OJ. 2010 Herbivore physiological response to predation risk and implications for ecosystem nutrient dynamics. Proc. Natl. Acad. Sci. 107, 15503–15507. (doi:10.1073/pnas.1009300107)

32. Shamir Weller ND, Raubenheimer D, Hawlena D. 2024 Constraints and demands interact to affect prey dietary reaction to predation. Funct. Ecol. 38, 2099–2109. (doi:10.1111/1365-2435.14647)

33. Cuff JP et al. 2021 MEDI: Macronutrient extraction and determination from invertebrates, a rapid, cheap and streamlined protocol. Methods Ecol Evol 12, 593–601. (doi:10.1111/2041-210X.13551)

34. Banerjee and Thaker. in press. A nutritionally-explicit test of the food-safety trade-off under predation risk. Scientific Reports

35. Batabyal A, Balakrishna S, Thaker M. 2017 A multivariate approach to understanding shifts in escape strategies of urban lizards. Behav. Ecol. Sociobiol. 71, 83. (doi:10.1007/s00265-017-2307-3)

36. Martín J, López P. 2004 Iberian rock lizards (*Lacerta monticola*) assess short-term changes in predation risk level when deciding refuge use. J. Comp. Psychol. 118, 280–286. (doi:10.1037/0735-7036.118.3.280)

37. Martín J, López P, Polo V. 2009 Temporal patterns of predation risk affect antipredator behaviour allocation by iberian rock lizards. Anim. Behav. 77, 1261–1266. (doi:10.1016/j.anbehav.2009.02.004)

38. Martin J. 2001 Repeated predatory attacks and multiple decisions to come out from a refuge in an alpine lizard. Behav. Ecol. 12, 386–389. (doi:10.1093/beheco/12.4.386)

39. Creel S. 2018 The control of risk hypothesis: Reactive vs. proactive antipredator responses and stress mediated vs. food mediated costs of response. Ecol. Lett. 21, 947– 956. (doi:10.1111/ele.12975)

40. Jimeno B, Hau M, Verhulst S. 2018 Corticosterone levels reflect variation in metabolic rate, independent of ‘stress’. Sci. Rep. 8, 13020. (doi:10.1038/s41598-018-31258-z)

41. Banerjee A, Fahis KT, Joshi M, Raubenheimer D, Thaker M. 2025 Does seasonal variation in the corticosterone response affect the nutritional ecology of a free ranging lizard? J. Anim. Ecol. 94, 627–641. (doi:10.1111/1365-2656.14249)

42. Clinchy M, Zanette L, Charlier TD, Newman AEM, Schmidt KL, Boonstra R, Soma KK. 2011 Multiple measures elucidate glucocorticoid responses to environmental variation in predation threat. Oecologia 166, 607–614. (doi:10.1007/s00442-011-1915-2)

43. Seebacher F, White CR, Franklin CE. 2015 Physiological plasticity increases resilience of ectothermic animals to climate change. *Nat*. Clim. Change 5, 61–66. (doi:10.1038/nclimate2457)

44. Jessop TS, Purwandana D, Imansyah MJ, Ciofi C, Jackson Benu Y, Arieifandy A. 2022 The influence of tropical seasonality on breeding phenology, growth, survival and movement of a large reptile (*Varanus komodoensis*). Biol. J. Linn. Soc. 136, 552–565. (doi:10.1093/biolinnean/blac045)

