## Supplementary image and tables for "Does diet influence physiological and behavioural responses to predation risk?"

**Supplementary Information**

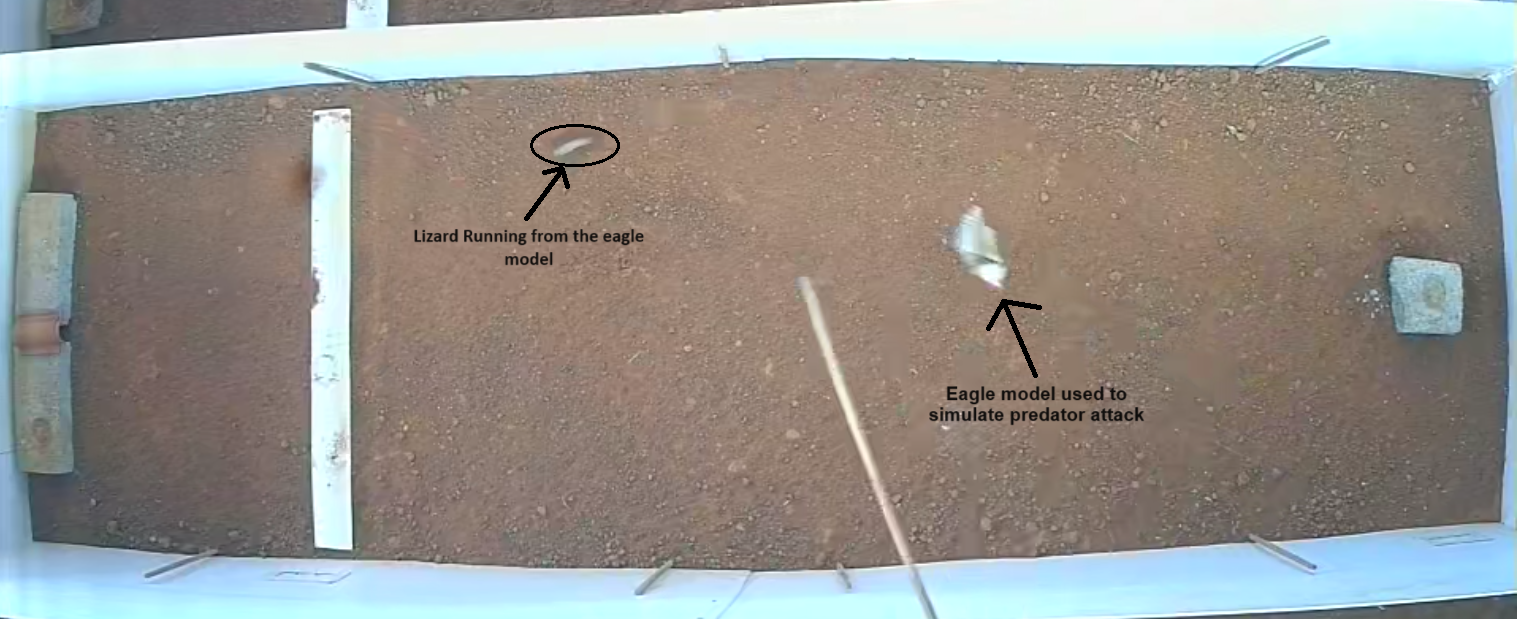

Figure S1: Snapshot taken during a simulated attack showing the bird model used to simulate attack and the focal lizard running towards the refuge.

Table S1: Output from the GLMM predicting the amount of mealworms (g) eaten (excluding zero values) as a function of Phase (safe vs predator exposure), Diet (high vs low NPE), and Phase:Diet, with Lizard ID as a random factor. Model family = Gamma (link = "log"). Dispersion (ϕ) = 0.102, The Variance and Std.Dev for Lizard ID were 0.034 and 0.183.

| Predictors | Estimate | Standard Error | Z value | p(>\|z\|) |
| --- | --- | --- | --- | --- |
| Intercept | 0.44 | 0.06 | 7.14 | <0.001 |
| Phase: Predator Exposure | -0.11 | 0.07 | -1.62 | 0.105 |
| Diet: Low NPE | -0.01 | 0.09 | -0.11 | 0.910 |
| Predator Exposure x Low NPE | 0.10 | 0.10 | 1.02 | 0.306 |

Table S2: Output from the GLMM predicting the proportion of time spent in refuge as a function of Phase (safe vs. predator exposure), Diet (high vs low NPE), and Phase:Diet, with Lizard ID as a random factor. Model family = Beta (link = "logit"). Dispersion (ϕ) = 10.2, The Variance and Std.Dev for Lizard ID were 0.40 and 0.62.

| Predictors | Estimate | Standard Error | Z value | p(>\|z\|) |
| --- | --- | --- | --- | --- |
| Intercept | -1.71 | 0.21 | -8.22 | <0.001 |
| Phase: Predator Exposure | 1.76 | 0.21 | 8.44 | **<0.001** |
| Diet: High NPE | -0.38 | 0.29 | -1.28 | 0.199 |
| Predator Exposure x High NPE | 0.23 | 0.29 | 0.79 | 0.431 |

Table S3: Output from the GLMM predicting the proportion of time spent on perch near the refuge as a function of Phase (safe vs. predator exposure), Diet (high vs low NPE), and Phase:Diet, with Lizard ID as a random factor. Model family = Beta (link = "logit"). Dispersion (ϕ) = 20.3, The Variance and Std.Dev for Lizard ID were 0.07 and 0.26.

| Predictors | Estimate | Standard Error | Z value | p(>\|z\|) |
| --- | --- | --- | --- | --- |
| Intercept | -0.73 | 0.11 | -6.61 | <0.001 |
| Phase: Predator Exposure | -0.87 | 0.15 | -5.67 | **<0.001** |
| Diet: High NPE | 0.03 | 0.15 | 0.21 | 0.835 |
| Predator Exposure x High NPE | 0.16 | 0.21 | 0.76 | 0.447 |

Table S4: Output from the GLMM predicting the proportion of time spent on perch far from the refuge as a function of Phase (safe vs. predator exposure), Diet (high vs low NPE), and Phase:Diet, with Lizard ID as a random factor. Model family = Beta (link = "logit"). Dispersion (ϕ) = 115, The Variance and Std.Dev for Lizard ID were 0.33 and 0.58.

| Predictors | Estimate | Standard Error | Z value | p(>\|z\|) |
| --- | --- | --- | --- | --- |
| Intercept | -3.58 | 0.17 | -21.57 | <0.001 |
| Phase: Predator Exposure | -0.49 | 0.18 | -2.80 | **0.005** |
| Diet: High NPE | 0.60 | 0.22 | 2.76 | **0.006** |
| Predator Exposure x High NPE | -0.54 | 0.23 | -2.37 | **0.017** |

Table S5: Odds ratios for the effects of Phase, Diet, and their interaction on the proportion of time spent on a perch far from the refuge. (Exponentiated from log odds).

| Contrast | | Estimate | Standard Error | Confidence Interval | z.ratio | p.value |
| --- | --- | --- | --- | --- | --- | --- |
| Predator Exposure | High NPE vs Low NPE | 1.07 | 0.25 | [0.56, 2.02] | 0.26 | 0.994 |
| Safe | High NPE vs Low NPE | 1.83 | 0.22 | [1.04, 3.21] | 2.76 | **0.030** |
| High NPE | Predator Exposure vs Safe | 0.36 | 0.15 | [0.24, 0.53] | -6.78 | **<0.001** |
| Low NPE | Predator Exposure vs Safe | 0.61 | 0.18 | [0.39, 0.96] | -2.79 | **0.027** |

Table S6: Output from the GLMM predicting the hiding duration after an attack as a function of Attack Number (1-6), Diet (high vs low NPE), and Attack Number:Diet, with Lizard ID as a random factor. Model family = Gamma (link = "log"). Dispersion (ϕ) = 0.30, The Variance and Std.Dev for Lizard ID was 0.18 and 0.43.

| Predictors | Estimate | Standard Error | Z value | p(>\|z\|) |
| --- | --- | --- | --- | --- |
| Intercept | 7.78 | 0.12 | 65.92 | <0.001 |
| Attack 2 | -0.14 | 0.11 | -1.24 | 0.213 |
| Attack 3 | -0.41 | 0.11 | -3.70 | **<0.001** |
| Attack 4 | -0.70 | 0.11 | -6.23 | **<0.001** |
| Attack 5 | -0.69 | 0.11 | -6.10 | **<0.001** |
| Attack 6 | -0.55 | 0.11 | -4.87 | **<0.001** |
| Diet: Low NPE | -0.003 | 0.17 | -0.02 | 0.986 |
| Attack 2 x Low NPE | -0.24 | 0.16 | -1.48 | 0.138 |
| Attack 3 x Low NPE | -0.23 | 0.16 | -1.44 | 0.150 |
| Attack 4 x Low NPE | -0.05 | 0.16 | -0.28 | 0.778 |
| Attack 5 x Low NPE | 0.07 | 0.16 | 0.44 | 0.659 |
| Attack 6 x Low NPE | -0.08 | 0.16 | -0.47 | 0.638 |

Table S7: Pairwise comparisons of hiding durations across attacks.

| Contrast | Estimate | Standard Error | z.ratio | p.value |
| --- | --- | --- | --- | --- |
| Attack 1 - Attack 2 | 0.26 | 0.08 | 3.22 | **0.016** |
| Attack 1 - Attack 3 | 0.53 | 0.08 | 6.59 | **<0.001** |
| Attack 1 - Attack 4 | 0.72 | 0.08 | 8.97 | **<0.001** |
| Attack 1 - Attack 5 | 0.65 | 0.08 | 8.07 | **<0.001** |
| Attack 1 - Attack 6 | 0.59 | 0.08 | 7.24 | **<0.001** |
| Attack 2 - Attack 3 | 0.27 | 0.08 | 3.38 | **0.009** |
| Attack 2 - Attack 4 | 0.47 | 0.08 | 5.79 | **<0.001** |
| Attack 2 - Attack 5 | 0.39 | 0.08 | 4.90 | **<0.001** |
| Attack 2 - Attack 6 | 0.33 | 0.08 | 4.08 | **<0.001** |
| Attack 3 - Attack 4 | 0.20 | 0.08 | 2.42 | 0.151 |
| Attack 3 - Attack 5 | 0.12 | 0.08 | 1.53 | 0.646 |
| Attack 3 - Attack 6 | 0.06 | 0.08 | 0.76 | 0.975 |
| Attack 4 - Attack 5 | -0.07 | 0.08 | -0.89 | 0.948 |
| Attack 4 - Attack 6 | -0.13 | 0.08 | -1.63 | 0.556 |
| Attack 5 - Attack 6 | -0.06 | 0.08 | -0.75 | 0.975 |

Table S8: Output from the GLM predicting the baseline corticosterone concentration as a function of Diet (high vs low NPE) and SVL (108-145 mm). Model family = Gamma(link = "log"). Dispersion (ϕ) = 0.29.

| Predictors | Estimate | Standard Error | Z value | p(>\|z\|) |
| --- | --- | --- | --- | --- |
| Intercept | 4.07 | 1.58 | 2.57 | 0.014 |
| Diet: Low NPE | 0.40 | 0.16 | 2.54 | **0.015** |
| Snout-Vent Length | 0.01 | 0.01 | 0.50 | 0.617 |
